# Title: Basal constriction during midbrain-hindbrain boundary morphogenesis is mediated by Wnt5b and Focal Adhesion Kinase

**DOI:** 10.1101/251132

**Authors:** Jennifer H. Gutzman, Ellie Graeden, Isabel Brachmann, Sayumi Yamazoe, James K. Chen, Hazel Sive

**Author notes:** equal contribution (Jennifer H. Gutzman, Ellie Graeden).

## Abstract

Basal constriction occurs at the zebrafish midbrain-hindbrain boundary constriction (MHBC) and is likely a widespread morphogenetic mechanism. 3D reconstruction demonstrates that MHBC cells are wedge-shaped, and initially constrict basally, with subsequent apical expansion. *wnt5b* is expressed in the MHB and is required for basal constriction. Consistent with a requirement for this pathway, expression of dominant negative Gsk3β overcomes *wnt5b* knockdown. Immunostaining identifies focal adhesion kinase (Fak) as active in the MHB region, and knockdown demonstrates Fak is a regulator of basal constriction. Tissue specific knockdown further indicates that Fak functions cell autonomously within the MHBC. Fak is epistatic to *wnt5b*, suggesting that Wnt5b signals locally as an early step in basal constriction and acts together with more widespread Fak activation. This study delineates signaling pathways that regulate basal constriction during brain morphogenesis.

## INTRODUCTION

Basal constriction is a cell shape change associated with zebrafish midbrain-hindbrain boundary constriction (MHBC) (Gutzman et al., 2008). This process is in contrast to the widely studied morphogenetic mechanism of apical constriction (Martin and Goldstein, 2014). Following our initial identification of the process, basal constriction has been described in several other systems and developmental processes. It is required for zebrafish and medaka optic cup morphogenesis (Bogdanovic et al., 2012; Martinez-Morales et al., 2009; Nicolas-Perez et al., 2016; Sidhaye and Norden, 2017), for notochord cell elongation in *Ciona*, (Dong et al., 2011), and for egg chamber elongation in *Drosophila* (He et al., 2010). Together these findings suggest that basal constriction is a conserved and fundamental morphogenetic process.

We previously demonstrated that basal constriction in the MHBC cells of the neuroepithelium requires an intact basement membrane and is laminin-dependent (Gutzman et al., 2008). During optic cup morphogenesis, basal constriction has been demonstrated to require actomyosin contraction and is also dependent on laminin (Nicolas-Perez et al., 2016; Sidhaye and Norden, 2017). However, the upstream signaling pathways that promote basal constriction have not been identified.

Since basal constriction at the MHBC occurs within a small group of cells, one hypothesis is that there is a localized signaling process involved, within the neuroepithelial sheet. Wnt-PCP signaling is one candidate regulatory pathway. Wnts are crucial for multiple morphogenetic events, including gastrulation, convergent extension, cell migration, and cell adhesion (Ciani and Salinas, 2005) and have been studied during development of the midbrain-hindbrain boundary (Buckles et al., 2004; Gibbs et al., 2017). Wnt5b, a known mediator of morphogenetic events in development is a regulator of cell shape and cell movement. It is required during gastrulation (Jopling and den Hertog, 2005; Kilian et al., 2003; Lin et al., 2010), mesenchymal cell migration and adhesion (Bradley and Drissi, 2011), *Xenopus* bottle cell apical constriction (Choi and Sokol, 2009), and tail morphogenesis (Marlow et al., 2004). In this communication, we demonstrate expression of *wnt5b* at the zebrafish MHBC and find a connection between Wnt5b, Gsk3β and Focal Adhesion Kinase, providing the first delineation of a signaling pathway required for basal constriction.

## RESULTS AND DISCUSSION

### Basally constricted cells are wedge-shaped

To delineate the steps in basal constriction, we examined cell shape during midbrain-hindbrain boundary constriction (MHBC) morphogenesis by injecting wild-type embryos with membrane targeted GFP (mGFP) and imaging using live confocal microscopy (Fig. 1A-D). Morphogenesis takes place beginning at approximately the 18 somite stage (ss) and extends to the prim-6 stage. At the start, the neural tube is composed of a pseudostratified epithelium with established apical-basal polarity (Fig. 1A). We identified three steps in MHBC morphogenesis. First, cells get shorter; second they form a basal constriction, and third they become apically expanded (Fig. 1A-H) (Gutzman et al., 2008; Gutzman et al., 2015). 3D reconstruction of MHBC cells revealed that as the cells basally constrict and apically expand they become wedge-shaped (Fig. 1C-D,G-H). The average basal anteroposterior width of the MHBC cells decreases from 2.1 microns to less than 0.5 microns between 14 ss, when the neuroepithelial cells are uniform in shape, and prim-6, when the cells have become wedge-shaped (Fig. 1I). The progression of MHBC cell shape changes is summarized in Fig. 1J-M. The ability to separate morphogenetic steps temporally supports recent data that discrete molecular and mechanical processes are likely to underlie each step. For example, during early MHB morphogenesis, differential roles were identified for non-muscle myosin IIA and non-muscle myosin IIB in mediating cell shortening and cell narrowing, respectively (Gutzman et al., 2015), and a role for calcium signaling was identified in regulation of cell length, and not cell width, specifically in MHBC cells (Sahu et al., 2017).

**Fig. 1.**
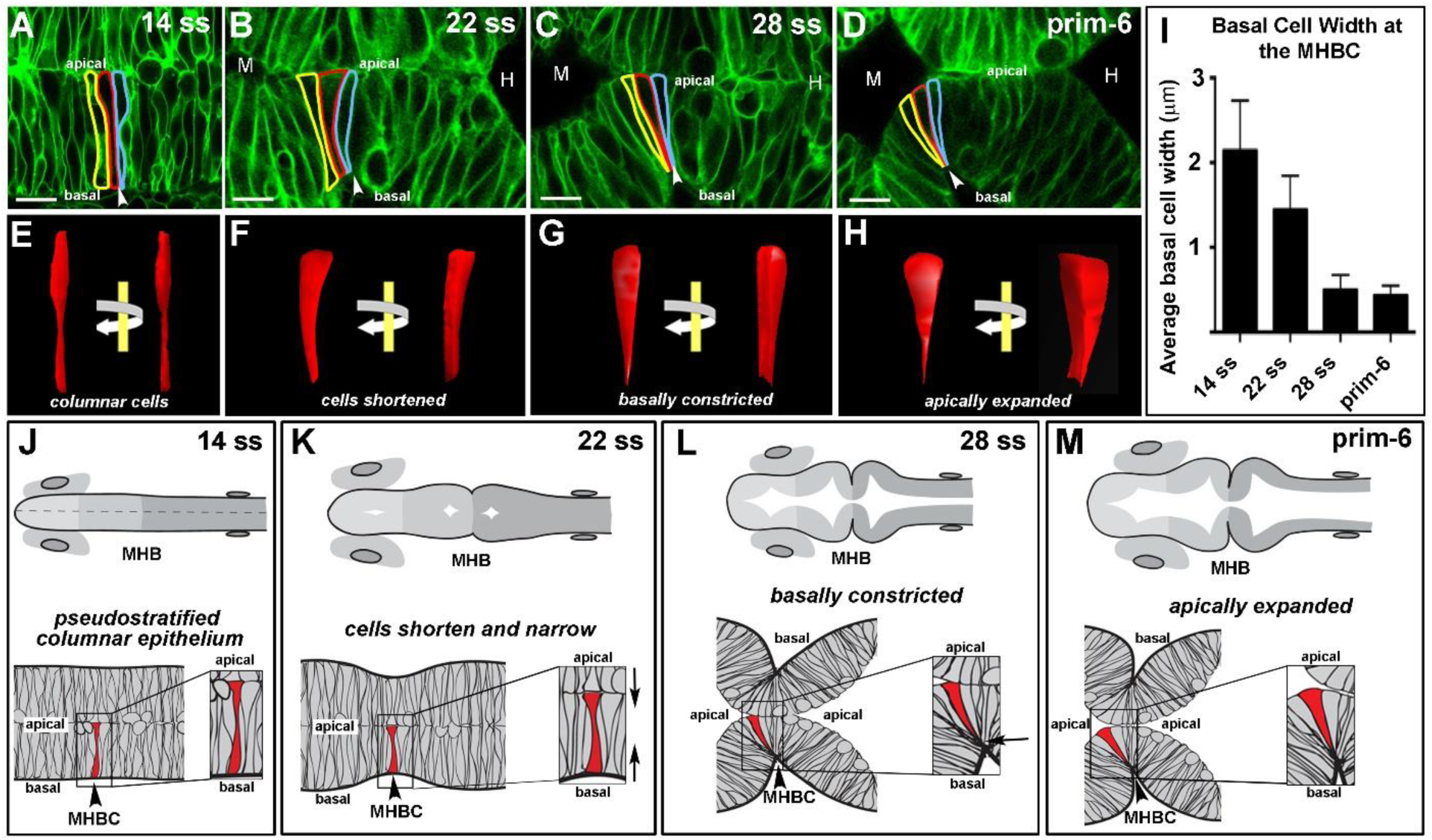
Basal constriction at the zebrafish MHBC occurs prior to apical expansion and results in wedge-shaped cells. (A-D) Live scanning confocal imaging of wild-type embryosinjected with mGFP mRNA and imaged at 14 ss, 22, ss, 28 ss, and prim-6. Cells at the MHBC are outlined in yellow, red, and blue. (E-H) 3D reconstruction of red outlined cell using 3D Doctor (Able Software). Each reconstruction is shown from two viewpoints. Face on as in the confocal image, and with a 45° rotation of the same image. (I) Histogram comparing the basal width of cells at each time point. (J-M) Schematics of wild-type MHBC formation. Anterior is to the left in all images. Arrowheads indicate the MHBC. M, midbrain; H, hindbrain. A-D, *n*>8 embryos per stage; I, *n=*3 embryos with 6 cells measured per embryo for each time point. Error bars reflect +/- s.d. Scale bars: A-D 9μm.

### wnt5b regulates basal constriction, possibly through Gsk3β

We hypothesized that genes required for basal constriction would be expressed prior to the start of MHBC formation and that expression would be restricted to cells undergoing basal constriction. In assessing the literature, we identified *wnt5b* expression as potentially correlating with MHBC morphogenesis both temporally and spatially (Montero-Balaguer et al., 2006; Thisse, 2005). We demonstrated that *wnt5b* RNA was enriched at the MHBC, using *in situ* hybridization (Fig. 2A-D). There is a low level of *wnt5b* expression throughout the embryo; however, expression increases at the MHBC shortly before morphogenesis begins and persists in this region throughout basal constriction (Fig. 2A-D). To determine the functional significance of Wnt5b in MHBC basal constriction, we used the established *wnt5b* antisense-morpholino modified oligonucleotide (MO) to inhibit function (Lele et al., 2001; Robu et al., 2007; Young et al., 2014). One-cell stage embryos were co-injected with control MO or *wnt5b* splice-site targeting MO with mGFP and live confocal imaging employed to examine cell shape. Consistent with expression, knockdown of *wnt5b* prevented basal constriction of cells at the MHBC (Fig. 2 E-F’). MHBC defects could be a result of anomalous patterning in the MHB, that occurs earlier during development. However, expression of the patterning genes *fgf8* and *pax2*, that are required for MHB formation, was unchanged in *wnt5b* morphants relative to control (Fig. S1). We also confirmed that *wnt5b* knockdown did not disrupt neuroepithelial cell apical-basal polarity (Fig. S1). These data suggest that requirement for *wnt5b* in basal constriction is not due to loss of early tissue patterning or polarity but is a later effect, perhaps directly impacting morphogenesis.

**Fig 2.**
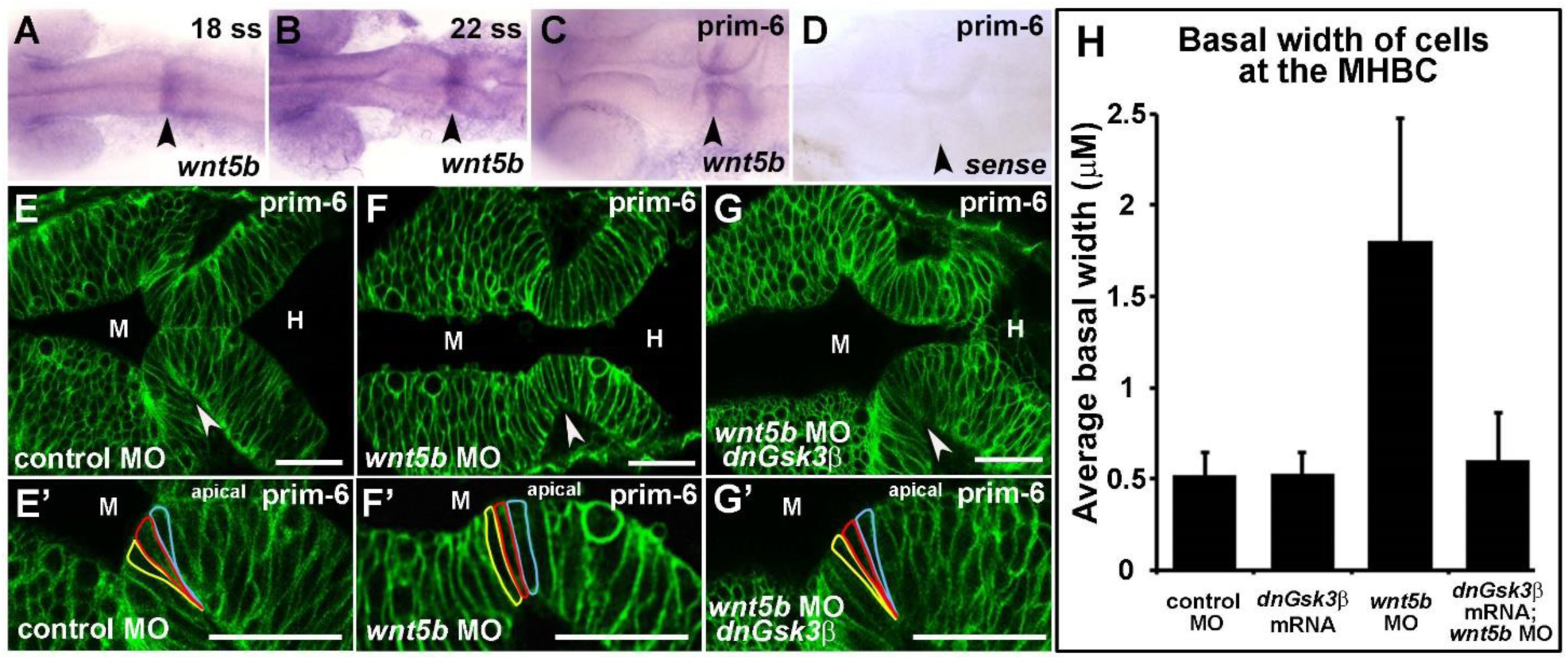
Wnt5b regulates basal constriction possibly through Gsk3β. (A-D) *In situ* hybridization of *wnt5b* expression during MHB development at 18 ss (A), 22 ss (B), and prim-6 (C). (D) prim-6 sense probe control. (E-G’) Live confocal images of the MHB region in prim-6 stage embryos. Single-cell wild-type embryos were injected with mGFP to label cell membranes and co-injected with control MO (E,E’), *wnt5b* MO (F, F’), or *wnt5b* MO and dnGsk3β mRNA (G,G’). (H) Quantification of basal cell width in control MO, *wnt5b* MO, *dnGsk3*β mRNA (image not shown), and *wnt5b* MO+*dnGsk3*β mRNA injected embryos. (H) For each treatment group, *n*=3 embryos. For each embryo, 6 cells located at the MHBC were measured, 3 cells on each side. Arrowheads indicate MHBC. M, midbrain; H, hindbrain. Scale bars: 26 μm.

Wnt5b is a ligand that can activate non-canonical Wnt signaling through Rho and JNK, and can also act through inactivation of Gsk3β (De Rienzo et al., 2011; Niehrs and Acebron, 2010; Terrand et al., 2009; Torii et al., 2008). In zebrafish, Wnt5b functions as a negative regulator of Wnt/b-catenin activity (Westfall et al., 2003) and studies in *Hydra* suggest that, during evagination, Wnt5b may also be involved with cross-talk between the canonical and non-canonical Wnt signaling pathways (Philipp et al., 2009). We asked whether inhibition of Gsk3β is required for basal constriction, using a kinase-dead Gsk3β (dnGsk3β) that is an established dominant negative construct (De Rienzo et al., 2011; Yost et al., 1996). Supporting a connection between Wnt5b and inhibition of Gsk3β, co-injection of dnGsk3β together with the *wnt5b* MO, prevented deficits in basal constriction seen after injection of *wnt5b* MO alone (Fig. 2G-H). Consistent with this, abnormalities in the gross morphology of the MHBC after injection of *wnt5b* MOs was prevented by expression of the dnGSK3β (Fig. S2). These data are consistent with a pathway in which Wnt5b regulates basal constriction through inhibition of Gsk3β.

### Fak is required at the MHBC for basal constriction

Basal constriction at the MHBC and in the optic cup both require laminin (Gutzman et al., 2008; Nicolas-Perez et al., 2016), a component of the underlying basement membrane, which interacts with integrins to regulate epithelial cell adhesion, migration, and differentiation (Yamada and Sekiguchi, 2015; Yurchenco, 2015). Focal adhesion kinase (Fak), a non-receptor tyrosine kinase, is a regulator of adhesion and cell migration that is activated through intracellular interactions with integrins (Parsons et al., 1994; Schaller, 2010). We therefore hypothesized that Fak plays a role in MHBC basal constriction. Since a primary mechanism for Fak activation is via autophosphorylation at Tyr397 (Schaller, 2010), we specifically hypothesized that autophosphorylated Fak^Y397^ would be localized to the MBHC. An antibody specific to Fak autophosphorylation site Y397 stained both the apical and basal surfaces in the neural tube at 18 ss, 24 ss, and prim-6 (Fig. 3A-D).

**Fig 3.**
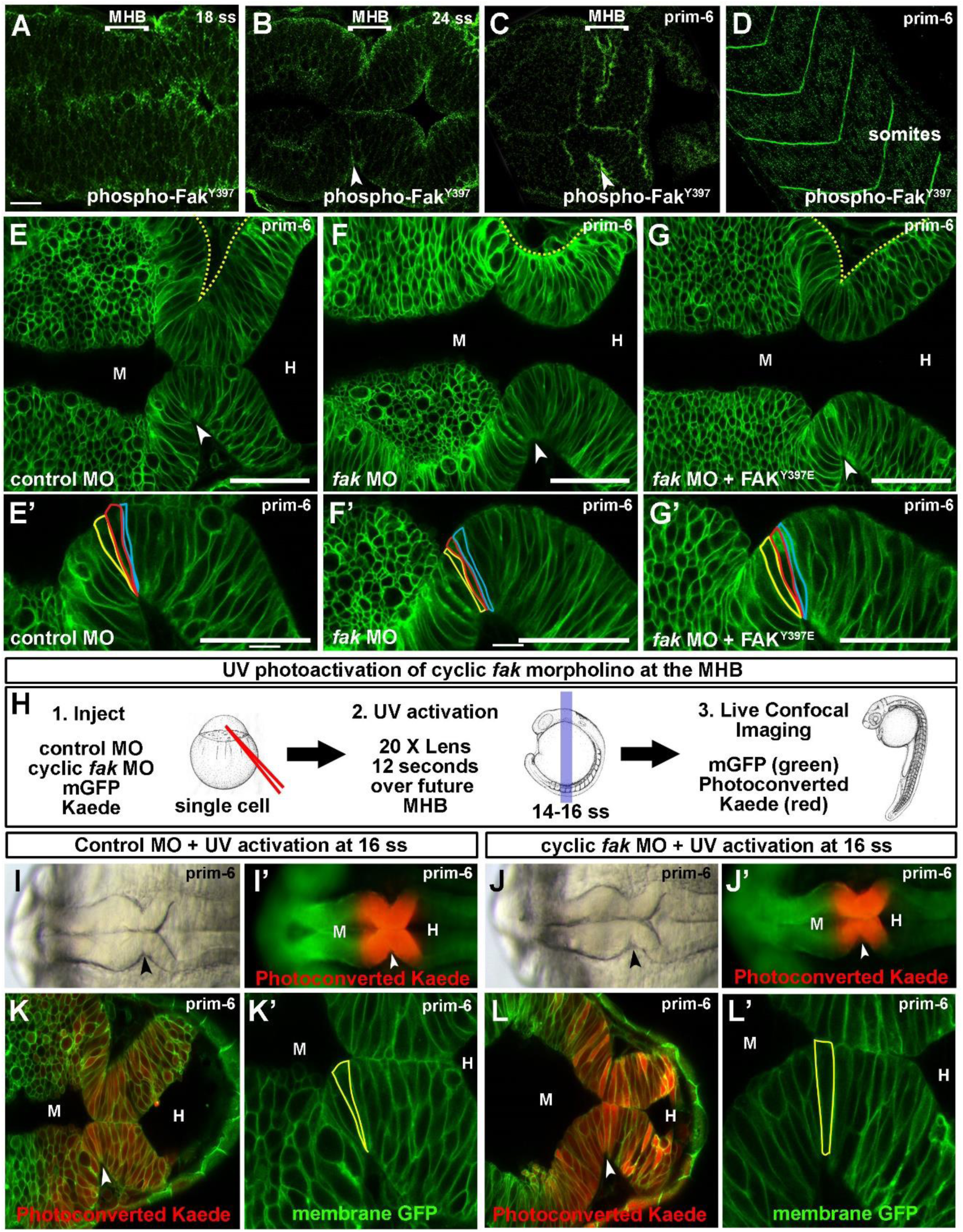
Focal Adhesion Kinase is required at the MHB for basal constriction. (A-D) Wildtype embryos stained with anti-phospho-Fak^Y397^ antibody and imaged by scanning confocal microscopy. (A,B) phospho-Fak^Y397^ is localized at the basal and apical sides of the neural tube at 18 and 24 ss. (C) Activated Fak is enriched at the MHBC at prim-6. (D) phospho-Fak^Y397^ is localized to somite boundaries at prim-6. (E-G’) Live confocal images of embryos co-injected with mGFP and control MO (E,E’), *fak* MO (F,F’), or *fak* MO + *FAK*^Y397E^ mRNA (G,G’). (E’-G’) Magnifications of individual cells outlined at the MHBC. (H) Schematic for *fak* caged MO experiments. 1. One-cell stage wild-type embryos were co-injected with mGFP and photoconvertable *Kaede* mRNA and either control MO or cyclic *fak* MO. 2. Cyclic *fak* MO was uncaged at 16 ss by UV activation. 3. Embryos were incubated until prim-6 and then imaged using brightfield, fluorescence, and live scanning confocal microscopy. (I-J’) Gross morphology using brightfield imaging (I,J) and corresponding fluorescent (I’,J’) images of embryos injected with control MO (I,I’) or photoactivatable *fak* MO (J,J’) after UV photoconversion. (K-L’) Live confocal images showing the MHB region of prim-6 embryos after photoconversion. (K’,L’) Magnifications of the neuroepithelium shown in K and L with individual cells outlined at the MHBC. (*n*>6). Anterior is to the left in all images. Arrowheads indicate MHBC. M, midbrain; H, hindbrain. Scale bars: A-C = 20μm. E-G’ = 50 μm.

We tested a role for Fak in basal constriction using knockdown with antisense-morpholino modified oligonucleotide injection. One-cell stage embryos were injected with control MO or a splice-site morpholino targeting the *ptk2.1* gene encoding one of the two *fak* genes in zebrafish. *fak* MO efficacy was confirmed by RT-PCR and Western blot analysis (Fig. S3A-C), and specificity was confirmed by rescue of *fak* MO injected embryos with co-injection of human FAK mRNA (Fig. S3). At the concentration used here, *fak* MO injections did not disrupt MHB tissue patterning or apical polarity markers (Fig. S1). *fak* morphants demonstrated disruption in MHB formation and abnormal basal constriction at the MHBC (Fig. 3E-F’). We tested activity of Fak^Y397^ in regulation of basal constriction using a phosphomimetic mutation of Tyr397 to Glu397 (Fak^Y397E^). Consistent with activation of Fak during basal constriction, co-injection of Fak^Y397E^ with *fak* MO was able to prevent abnormal basal constriction (Fig. 3G,G’).

We further tested the spatial and temporal requirement of Fak in the MHB to mediate basal constriction using injection of a photoactivatable cyclic *fak* MO. With UV activation, the cyclic *fak* MO becomes linear and binds to its target site (Yamazoe et al., 2012). We injected wild-type embryos with cyclic *fak* MO, Kaede mRNA, and mGFP mRNA. UV activation was performed at 14-16 ss, just before MHBC morphogenesis begins, in the MHB region as delineated by the change of Kaede from green to red. Basal constriction was disrupted after photoactivation of the cyclic *fak* MO, with no effect without activation or after UV treatment of the control MO-injected animals. These data show that Fak is required in the MHB region to mediate basal constriction (Fig. 3 G-K’). We further determined that Fak functions cell autonomously using MHB targeted cell transplantation (Fig. S4). Together these data indicate that Fak activity is required for basal constriction, and that Fak functions in the cells of the region that is undergoing basal constriction, beginning just prior to the start of the process.

### Wnt5b signals through Fak to mediate MHBC basal constriction

Since both Wnt5b and Fak are required for basal constriction, we asked whether there was an epistatic connection between the two signaling factors. We tested whether human FAK mRNA encoding the activated FAK^Y397E^ was able to prevent the basal constriction defect seen after Wnt5b inhibition. Indeed, we found that mRNA was able to rescue basal constriction in *wnt5b* morphants (Fig. 4A-G). This effect was not general, as FAK^Y397E^ did not rescue basal constriction defects found in laminin mutants ((Gutzman et al., 2008) and Fig. S5). These data indicate that Fak is epistatic to Wnt5b in activation of basal constriction at the MHBC.

**Fig 4.**
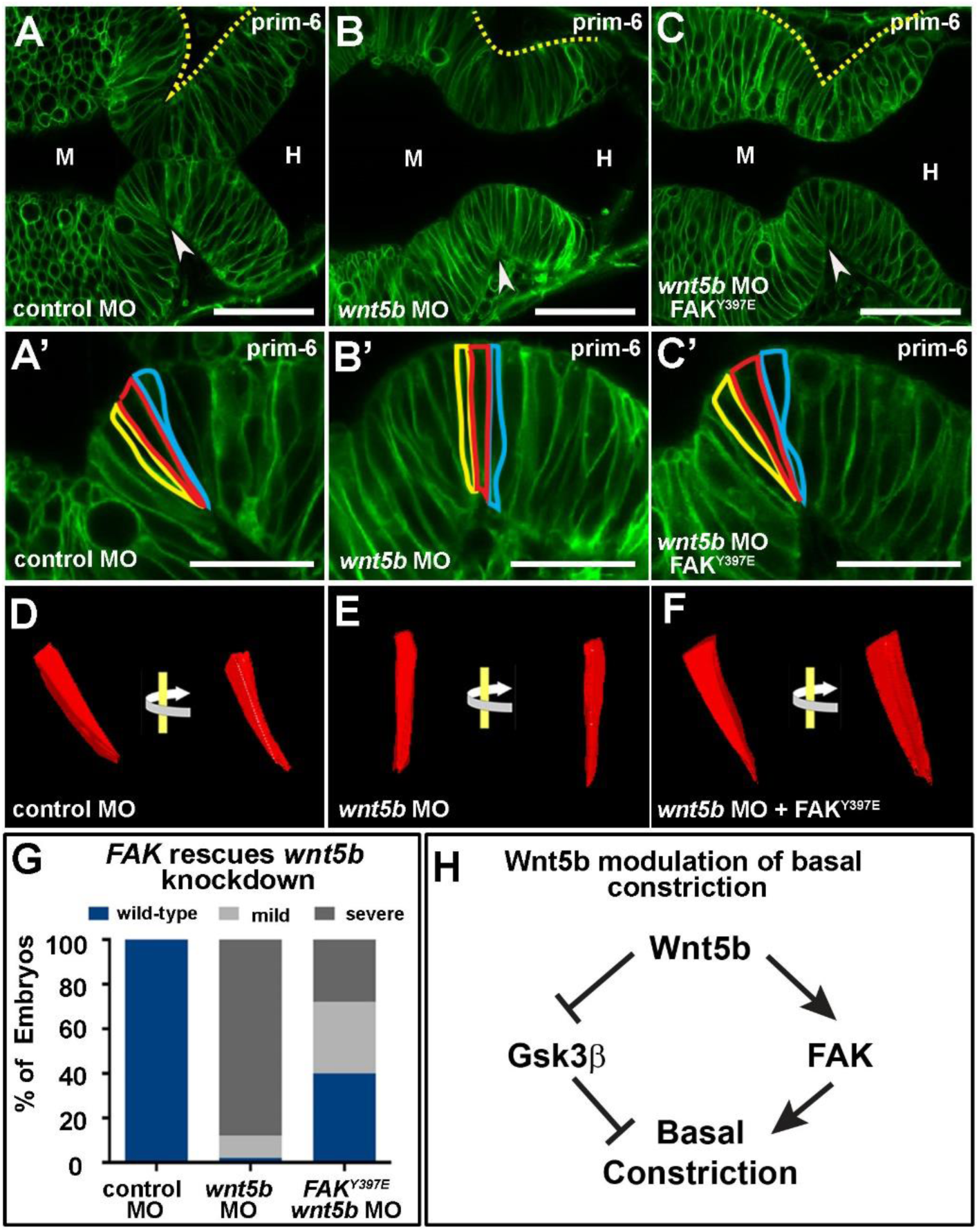
Focal Adhesion Kinase rescues effects of wnt5b knockdown on basal constriction. (A-C’) Live confocal images of the MHB region in prim-6 stage embryos. Single-cell wild-type embryos were co-injected with mGFP and control MO (A,A’), *wnt5b* MO (B, B’), or *wnt5b* MO and *FAK*^Y397E^ mRNA (C,C’). (D-F) 3D reconstruction of cells outlined in (A’-C’) using 3D doctor with view of the same cell rotated 45°. (G) Quantification of MHBC gross morphology and basal constriction following FAK^397E^ mRNA rescue of *wnt5b* knockdown (*n*>60 per condition). (H) Model pathway for Wnt5b regulation of basal constriction. Arrowheads indicate MHBC. M, midbrain. H, hindbrain. Scale bars: A-C, 50 μm; A’-C’ 25 μm.

Together, our results uncover key signaling factors contributing to basal constriction during MHBC morphogenesis. Our data point to a model in which Wnt5b signals locally at the MHBC as an early step in basal constriction, and acts together with more widespread Fak activation (Fig. 4H). We do not know whether or not Wnt5b and Fak are functioning at the same time during this process, nor whether their activity is also necessary for the earlier cell shortening or the subsequent apical expansion. Future experiments will uncover the molecular details of this signaling interaction and the role in other steps of MHBC formation.

## MATERIALS AND METHODS

### Zebrafish husbandry

Zebrafish lines were maintained, and embryo stages were determined, as previously described (Kimmel et al., 1995; Westerfield, 2000). Zebrafish strains used include wild-type AB and *sly*^m86^ (Schier et al., 1996).

### mRNA injections

All mRNA was *in vitro* transcribed with the mMessage mMachine kit (Thermo Fisher Scientific). Membrane-bound GFP (mGFP) mRNA was injected at 100-200 pg/embryo (kindly provided by B. Green, Dana-Farber Cancer Institute Boston, MA). Membrane-bound Cherry (mCherry) mRNA was injected at 50 pg/embryo (kindly provided by Dr. Roger Tsien, University of California San Diego). Photoconvertible Kaede mRNA was injected at 100 pg/embryo. pCS2+Kaede was kindly provided by Atsushi Miyawaki (RIKEN) (Ando et al., 2002). Human Focal Adhesion Kinase (FAK) (accession number BC035404) was purchased from Open Biosystems, EHS1001-5481173) and constructs were generated by subcloning into the pCS2+ expression plasmid. mRNA was *in vitro* transcribed and injected at 200-250 pg/embryo. pCS2+FAK was used as the backbone to generate the FAK^Y397E^ phosphomimetic using QuickChange Site-Directed Mutagenesis (Agilent). For each mRNA injection and rescue experiment, all embryos were injected with equal amounts of total mRNA. This included total mGFP when needed for imaging by scanning confocal microscopy. All microinjection experiments were performed at least three times.

### Live imaging and cell shape analysis

Live imaging of whole embryos was conducted using brightfield and fluorescent microscopy (SteREO Disvovery.V8, Zeiss). Live scanning confocal imaging was conducted as previously described (Graeden and Sive, 2009). Briefly, embryos were mounted inverted in 0.7% agarose and imaged using a 40X water immersion lens. Imaging was conducted using a Zeiss LSM510 or LSM720 scanning confocal microscope. Data was analyzed using Photoshop (Adobe) and Illustrator (Adobe) for cell outlines. 3D cell reconstruction was performed using 3D Doctor (Able Software). Individual cells at the MHBC were manually outlined in each *z*-section and rendered in 3D. A minimum of 6 embryos were imaged by scanning confocal microscopy and analyzed for basal constriction for each condition. Quantification of cell width was conducted using Imaris (Bitplane). The width of six cells at the MHBC from each of three embryos was measured at 300X zoom. Measurements were averaged and error bars reflect standard deviation for each condition.

### Morpholino injections

Splice site-blocking morpholino (MO) antisense oligonucleotides were injected into embryos at the one-cell stage. Morpholinos and concentrations used are as follows: 3 ng/embryo of *wnt5b* MO targeting the exon5/6 splice donor 5’-TGTTTATTTCCTCACCATTCTTCCG-3’ (Kim et al., 2005; Robu et al., 2007); 0.75 ng/embryo of *fak* MO (*ptk2.1*) targeting the exon 12/13 splice donor 5’-GTGTGTTTGGGTTCTCACCTTTCTG-3’; non-specific sequence standard control MO 5’-CCTCTTACCTCAGTTACAATTTATA-3’ at the concentration equal to the test condition; and *p53* MO 5’-GCGCCATTGCTTTGCAAGAATTG-3’ was co-injected at a concentration equal to 1.5 times the concentration of the test condition. Morpholinos were purchased from Gene Tools, LLC.

### Region specific knockdown by morpholino photoconversion

For photoactivatable morpholino experiments, we injected one-cell stage embryos with 1 ng/embryo of splice site-targeting cyclic *fak* MO (*ptk2.1*) 5’- GTGGGTGCTAACTGTCCGTCATATT-3’. The *fak* MO was cyclized with a photocleavable linker as previously described (Yamazoe et al., 2012) and remains inactive until “uncaging” by UV light. Linker photolysis reverts the MO to an linear oligonucleotide that can target the *fak* splice site. Embryos were co-injected with mGFP mRNA and Kaede mRNA together with either cyclic *fak* MO or control MO at the one-cell stage. Region and time specific UV activation was conducted at the 10-16 somite stage on cells located in the prospective MHB using a Zeiss Axioplan2 fluorescent microscope with a UV filter and adjustable iris. The tissue region that was activated by UV light is visible with the Kaede color change from green to red. Only cells that were photoconverted at the MHBC were analyzed for basal constriction as described.

### *In situ* hybridization

Antisense and sense RNA probes containing digoxigenin-11-UTP were synthesized from linearized plasmid DNA for *wnt5b* was obtained from Addgene #21282 (Stoick-Cooper et al., 2007). Standard methods for hybridization and for single color labeling were used as described (Sagerstrom et al., 1996). After staining, embryos were de-yolked, flat-mounted in glycerol and imaged with a Nikon compound microscope or a Zeiss Discovery V8.

### Fak^Y397^ immunostaining

Embryos were fixed in 4% paraformaldehyde; blocked in 2% normal goat serum, 1% BSA, and 0.1% Triton-X100 in PBT; incubated overnight at 4°C in primary antibody (anti-phosphoY397-FAK, 44-624 BioSource, Life Technologies), 1:200; then incubated in secondary antibody (goat anti-rabbit IgG conjugated with Alexa Fluor 488, Invitrogen, 1:500). Embryos were deyolked and mounted in glycerol. Images were collected using scanning confocal microscopy (Zeiss LSM510 or 710) and analyzed using Photoshop (Adobe).

## ACKNOWLEDGMENTS

We thank our Sive lab colleagues for useful criticism and Oliver Paugois for excellent fish care. We would like to acknowledge the W. M. Keck Foundation Biological Imaging Facility at the Whitehead Institute for advice and imaging assistance. pCS2+Kaede was kindly provided by Atsushi Miyawaki (RIKEN). This material is based upon work supported by the National Science Foundation Graduate Research Fellowship under Grant No. 1122374 to E.G.; National Science Foundation Grant no. 1258087 to H.S.; National Institutes of Health R01 GM087292 to J.K.C.; Massachusetts Institute of Technology CSBi/Merck Postdoctoral Fellowship to J.H.G; and Japan Society for the Promotion of Science Fellowship to S.Y.

